# Hyperoxia inhibits proliferation of retinal endothelial cells in Myc dependent manner

**DOI:** 10.1101/2020.11.09.375220

**Authors:** Charandeep Singh, Andrew Benos, Allison Grenell, Sujata Rao, Bela Anand-Apte, Jonathan E. Sears

## Abstract

Oxygen supplementation is necessary to prevent mortality of severely premature infants. However, the supraphysiological concentration of oxygen utilized in these infants simultaneously creates retinovascular growth attenuation and vasoobliteration that induces retinopathy of prematurity. Here, we report that hyperoxia regulates the cell cycle and retinal endothelial cell proliferation in a previously unknown Myc dependent manner which contributes to oxygen-induced retinopathy.

## Introduction

Retinopathy of prematurity (ROP) is a leading cause of infant blindness world-wide, accounting for 184,700 new cases annually (Blencowe et al., 2013; Hoppe et al., 2016; Sears et al., 2008). Although oxygen supplementation is necessary to prevent mortality in premature infants, oxygen supplementation in severely low birthweight infants can be detrimental to the developing premature organs, such as the retina, brain, and lung. Although ROP does not develop until corrected gestational age of 30-32 weeks, it is retinovascular growth attenuation and vasoobliteration caused by higher than *in utero* oxygen concentrations that creates increased avascular retinal tissue that causes pathological angiogenesis followed by retinal detachment and blindness (Kim et al., 2018). One of the early clinical signs of ROP is retinovascular growth suppression (Chen and Smith, 2007; Hartnett and Penn, 2012; Narayanan et al., 2014). A similar phenotype can be recapitulated in the mouse and rat model of oxygen induced retinopathy (OIR) (Barnett et al., 2010; Smith et al., 1994). This phenotype is often referred to as “oxygen toxicity” as it bears the negative connotation reflecting ill effects of oxygen on the vascular development. In vitro and in vivo studies have demonstrated that hyperoxia increases the formation of reactive oxygen and nitrogen species (Auten and Davis, 2009; Zou et al., 2019). Furthermore, hyperoxia upregulates neuronal apoptosis in the brain and the retina (Felderhoff-Mueser et al., 2004; Ikonomidou, 2009; Taglialatela et al., 1998; Terraneo et al., 2017; Yiş et al., 2008). Although neurons are non-mitotic fully differentiated cells, they do harbor cell cycle proteins and recent studies have demonstrated that dysregulation in cell cycle protein levels in neurons can lead to apoptosis. However, mitotic cells of non-neuronal origin can enter into a long G_0_ phase under unsuitable circumstances and can re-enter the cell cycle when conditions become favorable (Foster et al., 2010; Linke et al., 1996). In mice, hyperoxia affects the vasculature in early postnatal stages when the endothelial cells are still proliferating and migrating. Smith et. al. (1993) demonstrated that once the vasculature is fully developed, these mice do not develop the vaso-obliteration and neovascularization phenotype after exposer to 5 days of hyperoxia (Smith et al., 1994). This implies that susceptibility to hyperoxia is not merely caused by oxidative damage but involves more complex molecular pathways that are active in the early stages of retinal development. Like in retinal tissue, postnatal oxygen rich environment inhibits proliferation of cardiomyocytes (Puente et al., 2014). Mammalian cardiomyocytes have regenerative capacity at birth but lose this potential postnatally as the oxygen rich environment prevents cell proliferation. In mice, after postnatal day 7, cardiomyocytes become binucleated and permanently exit the cell cycle through DNA damage-induced cell cycle arrest (Puente et al., 2014). These differences in response to hyperoxia amongst cell types demonstrate the heterogeneity of cell cycle control and warrant closer examination of cell type-specific mechanisms.

Myc is a critical regulator of the cell cycle and cellular proliferation (Bretones et al., 2015b). Hypoxia-induced increase in HIF1 levels result in decreased Myc RNA and protein expression (Okuyama et al., 2010; Sun and Denko, 2014; Wise et al., 2011). Furthermore, Myc levels are inversely proportional to nutrient availability and cell density. Myc is downregulated during starvation conditions, halting the cell cycle, which leads to the loss of proliferation to protect the essential supplies for survival (Bretones et al., 2015b). One of the mechanisms by which Myc regulates cellular proliferation is via upregulating polyamine production (Bachmann and Geerts, 2018). The polyamine pathway is indispensable for normal proliferation and growth (Li et al., 1999; Tabor and Tabor, 1984). Hypoxia increases glycolysis by upregulating pyruvate dehydrogenase kinase-1 (PDK1) which phosphorylates pyruvate dehydrogenase (PDH) and thereby inhibits entry of glycolytic carbon into the TCA cycle. This switch in metabolic flux downregulates cell proliferation by inducing the expression of cyclin-dependent kinase inhibitor (CDKI). Although phosphorylation of PDH is HIF dependent, upregulation or downregulation of Myc by HIF or vice-versa is context dependent, and there have been no studies on effects of hyperoxia on Myc protein levels. HIF can suppress cell proliferation by inhibiting the transcriptional activity of Myc (by destabilizing Myc’s interaction with other transcriptional co-factors). Recent reports have shown that HIF1 displaces Myc from MYC-associated protein X (MAX), resulting in destabilization of Myc (Eilers and Eisenman, 2008; Grinberg et al., 2004). These findings appear to contradict the phenotype of OIR; if hyperoxia downregulates HIF, one might assume that Myc would be induced by hyperoxia. In this investigation, we analyzed the effect of hyperoxia on key cell cycle regulators. Our findings indicate a central effect of hyperoxia on Myc protein levels, providing a molecular mechanism of how oxygen induces cell cycle arrest in retinal endothelial cells.

## Results

To study the effect of hyperoxia on retinal endothelial cell proliferation, we cultured primary human retinal endothelial cells for 24 h under normoxic conditions, followed by hyperoxic or normoxic conditions for 4-6 days. Cellular proliferation was significantly reduced under hyperoxic conditions (Fig. 1a), despite the presence of mitogens such as VEGF, IGF and EGF (please refer to the materials and methods section for the complete media recipe).

**Figure 1.**
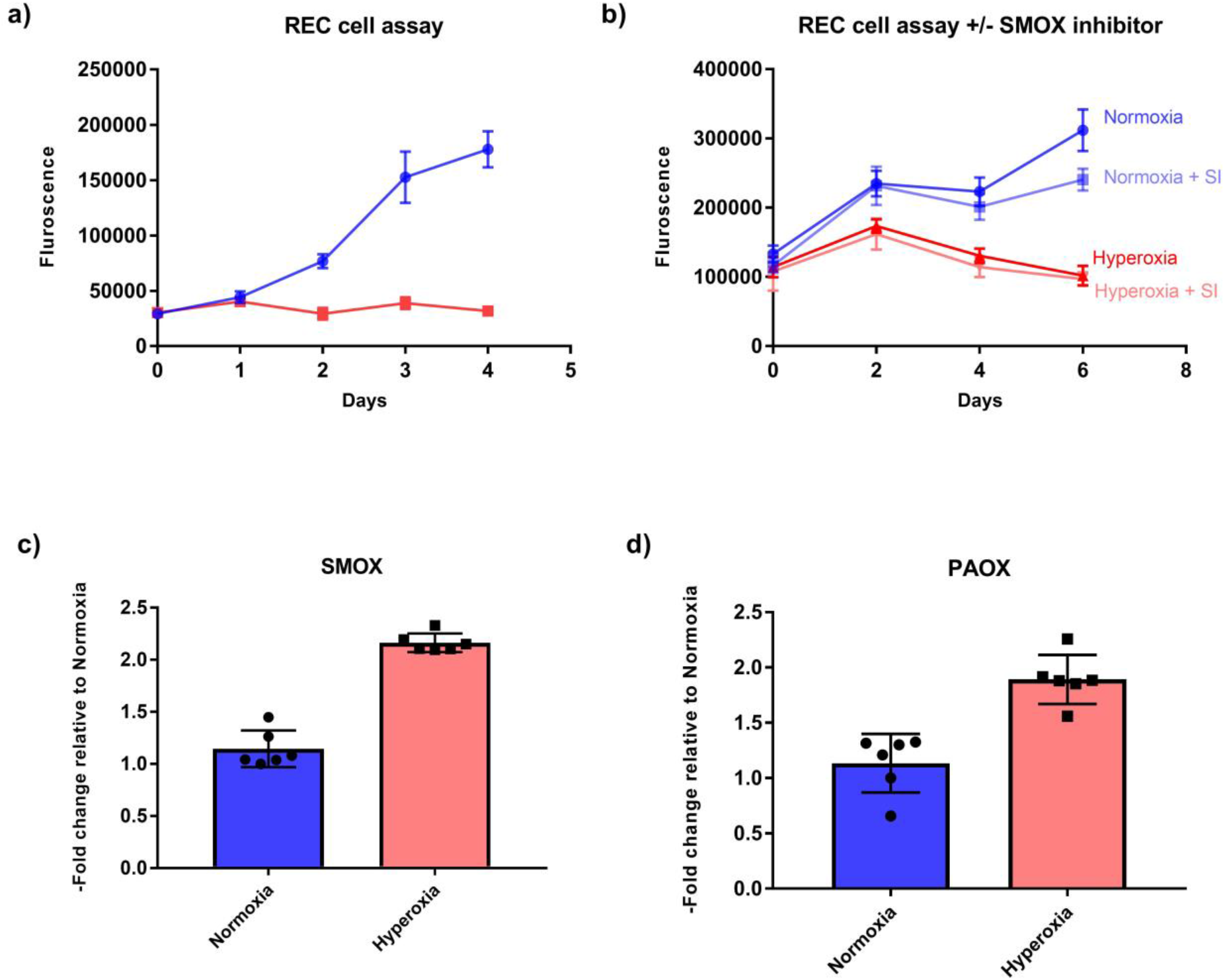
Hyperoxia inhibits prolifeartion and increases expression of polyamine oxidation genes in retinal endothelial cells. a) Retinal endothelial cells cultured in hyperoxia demonstrated proliferation defects even in the presence of growth factors (n=8 biological replicates per condition). b) Proliferation defects were not rescued by spermidine oxidase inhibitor (SI) MDL72527 (n=6 biological replicates per condition) c) Spermine oxidase (SMOX) t-test p-value <0.05 d) Peroxisomal N (1)-acetyl-spermine/spermidine oxidase (PAOX) t-test p-value <0.05 (n=6 biological replicates per condition).

We next examined the expression of polyamine oxidation/breakdown genes, as polyamine levels are critical regulators of cell proliferation in prokaryotes and eukaryotes (Igarashi and Kashiwagi, 2000). Polyamines modulate translation by making complexes with RNA. Critical enzymes in the polyamine pathway, such as ornithine decarboxylase (ODC), peak at G1/S and G2/M transition points, implying that polyamine levels control these checkpoints (Yamashita et al., 2013). In addition, Nakayama and Nakayama (1998) demonstrated that the cell cycle inhibitors p27Kip1 and p21Cip1/WAF1 were upregulated in response to low polyamine concentration in the cells (Nakayama and Nakayama, 1998). This finding was confirmed by Yamashita et al. (2013), who demonstrated that p27Kip1 translation was enhanced by polyamine deficiency (Yamashita et al., 2013). In the retina, oxidation/breakdown of polyamines increases in response to hyperoxia and induces neuronal death (Narayanan et al., 2014). Spermine oxidase (SMOX), an enzyme that catabolizes early substrates of the growth-inducing polyamine pathway, is reported to be increased in hyperoxic conditions (Narayanan et al., 2014). We investigated whether the expression levels of polyamine oxidation genes are regulated at transcriptional levels in response to hyperoxia. Hyperoxia indeed results in upregulation of SMOX in the endothelial cells as compared to normoxia (Fig. 1c). We also measured expression of another gene responsible for polyamine oxidation, Peroxisomal N (1)-acetyl-spermine/spermidine oxidase (PAOX) and found increased expression in response to hyperoxia (Fig. 1d). This implies that the polyamine oxidation genes are transcriptionally controlled in hyperoxic conditions. The SMOX inhibitor MDL 72527 has been shown to reduce retinal neuronal death in the OIR model (Narayanan et al., 2014). We determined that SMOX inhibition could not rescue the cell proliferation phenotype in endothelial cells cultured under hyperoxic conditions (Fig. 1b).

Given that the inhibition of enzymes that downregulate critical polyamines necessary for growth did not rescue the growth of hyperoxic endothelial cells, we further examined upstream cell cycle regulators in synchronized primary human retinal endothelial cells. Since Myc protein levels positively correlate with cell proliferation in many different cell types, we measured Myc protein levels in normoxic vs. hyperoxic endothelial cells. Myc protein levels were significantly reduced in hyperoxic endothelial cells compared to normoxic conditions (Fig. 2a). We confirmed Myc levels with an additional antibody (Fig. S1). However, we did not observe any changes in the Myc gene expression between normoxia and hyperoxia, implying Myc levels may be controlled by a previously unknown post-translational modification of Myc protein (Fig. S2). To further confirm the relationship between hyperoxia and cell cycle arrest, we next evaluated p53, because this established regulator of the cell cycle is reported to regulate the phosphorylation of Rb (pRb) (Kastenhuber and Lowe, 2017). p53 can either activate cell cycle arrest by inducing p21/Rb axis or apoptosis by inducing BCL-2 pathway. Despite this duality of function, it is reported that only one of these pathways is activated at a time; however it is not clear which cellular events determine which of these pathways could be activated (Hafner et al., 2019; Kastenhuber and Lowe, 2017). We measured p53 and p21 levels in normoxic and hyperoxic conditions. p53 (Fig. 2B,G) and p21 (Fig. 2D,I) levels were increased in hyperoxic conditions confirming cell cycle arrest. Taken together, these findings establish that hyperoxia causes cell cycle arrest in G1 phase, via p53 and Myc dependent pathways.

**Figure 2.**
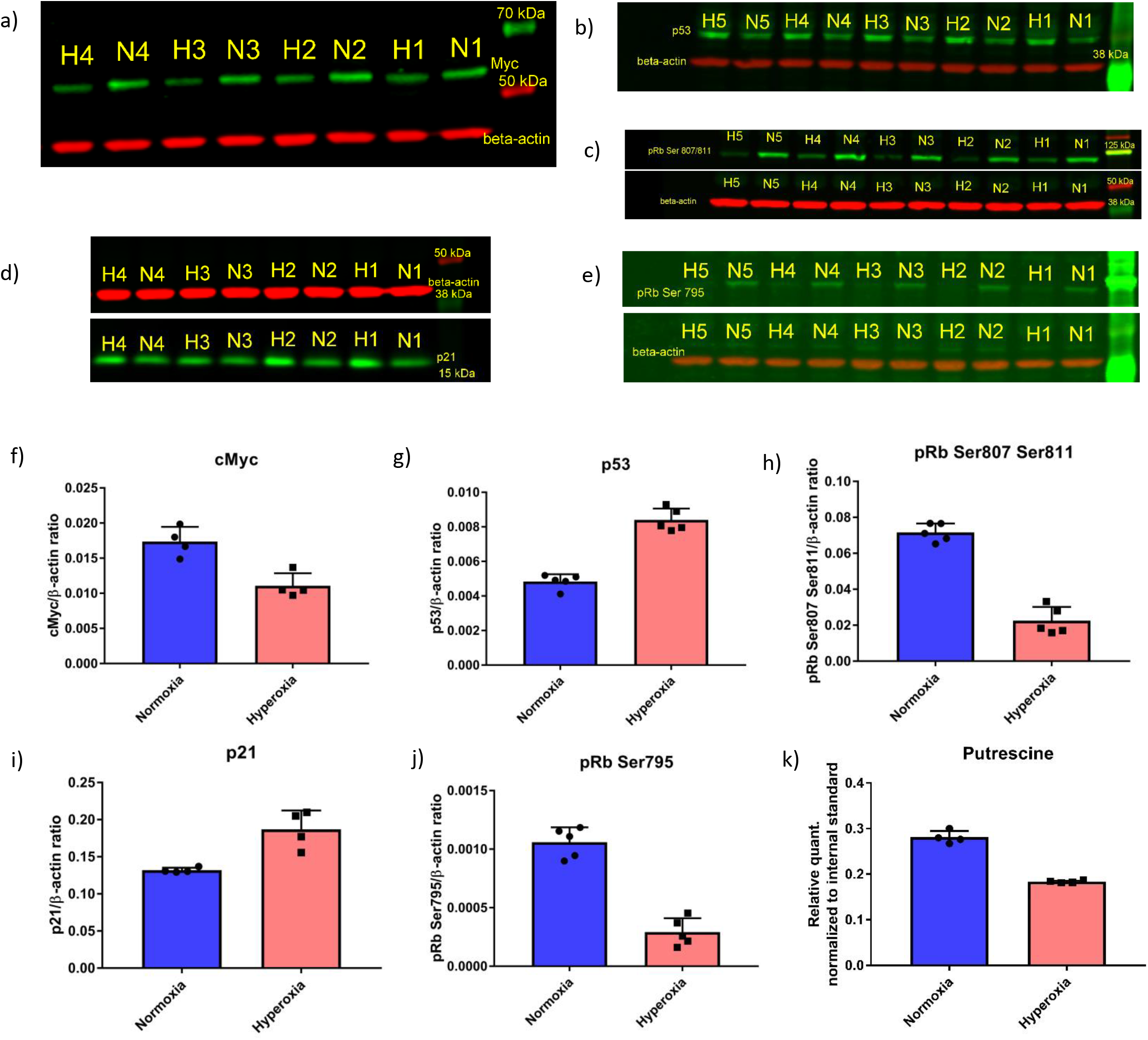
Cell cycle signaling proteins and polyamine (putrescine) affected by hyperoxia in endothelial cells. Western blots of normoxic (N1-5; each number represent biological replicate) and hyperoxic (H1-5; each number represent biological replicate) samples for a) Myc b) p53 c) pRb Ser 807/811 d) p21 e) pRb Ser 795. Quantification of the western blot is provided in the histograms f) Myc, g) p53, h) pRb Ser 807/811 i) p21 j) pRb Ser 795 k) Putrescine. t-test p-values for all the quantifications were all less than 0.05.

To further confirm that hyperoxia induces cell cycle arrest, we evaluated the phosphorylation of Retinoblastoma protein (Rb). The first step in committing cells to cell division is transition from the G1 to the S phase, which is dependent on phosphorylation of Retinoblastoma (Rb) protein (Bretones et al., 2015b; Knudsen and Wang, 1997). Phosphorylated Rb leads to the increased concentration of E2F elongation factor thereby signaling translation of the proteins required for S-phase (Bretones et al., 2015b; Knudsen and Wang, 1997). There are 19 known phosphorylation sites on human Rb1 protein (source: Uniport)(Consortium, 2018), of these the three most important sites involved in cell cycle regulation are Ser 807, Ser 811, and Ser 795 (Rubin et al., 2005). Recent work by multiple teams have highlighted that out of these three sites, Ser 807 and Ser 811 regulate c-Abl binding of Rb1 (Knudsen and Wang, 1997; Rubin et al., 2005). Ser795 is involved in binding of Rb1 to E2F transcription factor (Knudsen and Wang, 1997; Rubin et al., 2005). We measured the levels of phosphorylated pRb Ser807/811 (Fig.2C,H) and pRb Ser795 (Fig. 2E,J). Both the phosphorylated forms of Rb protein were decreased in response to hyperoxia, indicating cell cycle arrest in G1 phase. We additionally looked at the levels of putrescine and found it to be decreased in hyperoxic conditions (Fig. 2k). We confirmed these findings with an additional LC-MS/MS method (Fig. S3 and S4). Spermidine was not statistically significantly changed and spermine quantity was not high enough to be confidently measured.

## Discussion

Our results clearly demonstrate that hyperoxia downregulates endothelial cell proliferation, without inducing cell death, by decreasing expression of key cell cycle determinants. Although cell proliferation is controlled by multiple mechanisms under physiological conditions, p53 and Myc are reported to be the most important regulators of cell proliferation. Myc controls expression of positive regulators of cell cycle and also induces growth by down regulating the expression of cell cycle inhibitors such as p21CIP1/WAF1 (for review see Bretones, Delgado and Leon (2015))(Bretones et al., 2015a). The most widely accepted and recognized mechanism of p21 repression by Myc is through Miz-1. Miz-1, when in contact with Myc, represses p21. Miz-1/Myc interaction also makes p21 insensitive to p53 signaling (Peukert et al., 1997; Seoane et al., 2002). The significance of our observation of oxygen induced downregulation of Myc is that it demonstrates that the central paradigm of the inverse relationship of HIF and Myc expression may not hold true in hyperoxia. Downregulation of Myc in hyperoxia, when HIF1 levels are known to be decreased, is unexpected as Myc in most cases works antagonistically to HIF (Okuyama et al., 2010; Sun and Denko, 2014; Wise et al., 2011). This warrants further studies on how and why hyperoxia downregulates Myc and cell proliferation. Both Myc and p53 control these mechanisms in response to cellular stress like DNA damage or nutrient deprivation (Puente et al., 2014; Stine et al., 2015). Biomass synthesis pathways like serine/one-carbon and glutaminolysis involve Myc protein. These pathways were found to be altered by hyperoxia in our previous studies (Singh et al., 2019; Singh et al., 2018; Singh et al., 2020).

A second important finding from our experiments is that standard, HIF-induced mitogens are unable to override oxygen induced growth suppression, at least in retinal endothelial cells. VEGF and other mitogens are known to activate endothelial cell proliferation. In our experiments, hyperoxia was able to block cell proliferation of endothelial cells despite the presence of mitogens such as VEGF, IGF, and EGF – which implies that hyperoxia inhibits cell proliferation by acting downstream of these targets. Both MAPK and PI3K-Akt pathways control cell cycle progression and are downstream of VEGF and EGF/IGF. Our findings suggest the relevance of these downstream pathways to OIR. Another independent possibility is that the cells, in response to hyperoxia, have an aberrant VEFR2/R1 ratio rendering them less sensitive to mitogens.

In conclusion, our investigation demonstrates that hyperoxia downregulates retinal endothelial cell proliferation by downregulating Myc protein levels and upregulating p53 protein levels. The schema in Fig. 3 provides a summary of our findings and a potential blueprint for examining how hyperoxia induces retinal endothelial cell growth suppression.

**Figure 3.**
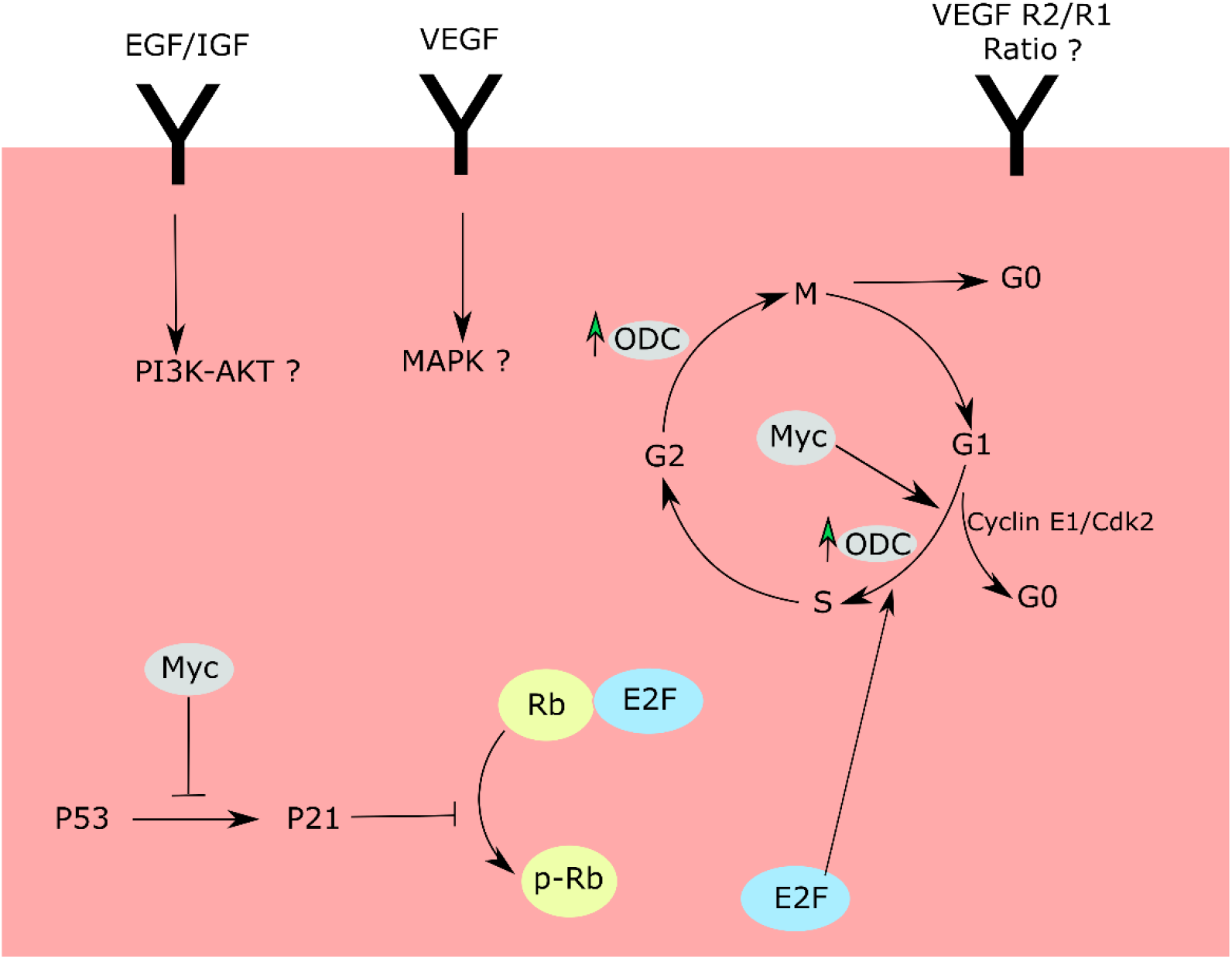
Cell cycle checkpoints and proposed routes affected by hyperoxia. Hyperoxia downregulates Myc and upregulates P53 proteins thereby increasing p21 protein levels. P21 further downregulates pRb levels, leading to cell cycle arrest in G1 phase. The cell cycle arrest can be due to anomalies in MAPK and P13K/Akt pathways downstream of VEGF, EFG/IGF receptors. Alternatively, the cell cycle changes can be a result of an aberrant VEGFR2/R1 ratio.

## Materials and methods

### Cell proliferation assay

Primary human retinal endothelial cells were purchased from Cell Systems and used within 4-5 passages. Cells were maintained in endothelial cell media from Cell Biologics (catalogue number H1168). Cells were plated in black 96-well plates overnight and then incubated in normoxic (21% oxygen) or hyperoxic (75% oxygen) incubator for 4-6 days. Cell proliferation was measured by using CyQuant™ NF cell proliferation assay kit from Invitrogen following protocol provided with the kit. SMOX inhibitor, MDL 72527, was spiked into the media at a final concentration of 100 μM.

### Protein extraction form cultured cells

Cells were plated in 100 mm x 20 mm dishes (Corning) coated with the attachment factor (Cell systems catalogue number 4Z0-201) and maintained in the media described above. Once the cells reached 70-80% confluency, plates were either incubated in either normoxic (21% oxygen) or hyperoxic (75% oxygen) incubators for the next 24 h. Both the incubators were set at 37 °C temperature and 5% CO_2_. After 24 h of exposure to different levels of oxygen, proteins were extracted from these cells. To extract the proteins, cells were briefly washed with normal saline, followed by addition of 300 μL of RIPA buffer containing cOmplete™ protease inhibitor and phosphatase inhibitor (both from Roche). Cells were scraped with cell scrapers and transferred to 1.5 tubes. Cells were briefly sonicated and then spun down in a centrifuge at 15000 x g for 15 min at 4°C. The supernatant was transferred to fresh tubes and stored at −80°C until further use.

### SDS-PAGE and western blotting

Protein concentration in the cell lysates was measured using BCA protein assay reagent (Pierce™). Protein sample 15-20 μg was mixed with tris-glycine SDS loading dye and 20 mM DTT. Samples were heated at 94°C for 3 min following centrifugation at 15000 x g at room temperature for 3 min. Supernatant 30 μL was loaded into each well of 4-20 % or 12% Tris-glycine Novex™ WedgeWell™ precast gel (Invitrogen). Equal quantities of protein samples were loaded in all the wells of each individual gel. Proteins were separated at constant voltage of 150 V. Proteins were transferred from gel to 0.45-micron PVDF membrane (Millipore) at 70 V for 2 h using wet-transfer in tris-glycine buffer. Following transfer, membranes were dried for 1 h then quickly rinsed with methanol followed by rinsing with water. Membranes were then washed with TBS and blocked with intercept TBS blocking buffer (LI-COR) for 1 h. Following blocking, membranes were treated with primary antibodies diluted in intercept TBS blocking buffer containing 0.2 % Tween 20 overnight at 4°C. Membranes were washed with TBST 3 times (5 minutes per wash) then treated with secondary antibodies diluted in intercept TBS blocking buffer containing 0.2 % Tween 20 and 0.01% SDS (w/v) solution, for 1 h at room temperature in the dark. Following incubation with secondary antibody, blots were washed 3 times with TBST and rinsed with TBS. Images were acquired on Odyssey® CLx imaging system (LI-COR). Images were analyzed using Image Studio Lite version 5.2 (LI-COR)

It has earlier been noted previously that the Myc antibodies binds to a non-specific band which co-elutes with endogenous Myc (Tibbitts et al., 2012). We also observed the non-specific band which eluted very closely with Myc. To circumvent this problem, we included an additional step of stripping and re-probing the blot for Myc protein. This step was necessary to remove a second non-specific band seen in our Myc blots. p21 western blot was stripped with 10 ml of Restore™ PLUS western blot stripping buffer (Thermo Fisher Scientific) for 20 min at room temperature followed by re-probing with Myc and β-actin antibody for 1h at room temperature. After treatment with primary antibody, above described procedure was used for secondary antibody treatment and imagining.

Following primary antibodies were used:

1. c-Myc (D84C12) Rabbit mAb catalog # 5605
2. Phospho-Rb (Ser 807/811) (D20B12) XP® Rabbit mAb catalog # 8516
3. Phospho-Rb (Ser 795) Rabbit antibody catalog # 9301
4. p21 Waf1/Cip1 (12D1) Rabbit mAb catalog # 2947
5. p53 Rabbit antibody catalog # 9282
6. ß-actin (8H10D10) Mouse mAb catalog # 3700

All the antibodies were purchased from Cell Signaling and were diluted as recommended by the vendor. Following secondary antibodies were used:

IRDye® 800CW Donkey (polyclonal) anti-Rabbit IgG (H+L), catalog number 925-32213 from LI-COR.

IRDye® 680RD Donkey (polyclonal) anti-mouse IgG (H+L), catalog number 925-68072 from LI-COR. Both the antibodies were used at 1:2000 dilution.

### RNA extraction and quantitative RT-PCR

Cells were cultured in 6-well plates and maintained in endothelial cell media in normoxic incubator. At around 70-80% confluence, cells were transferred to normoxic or hyperoxic incubator for 24 h, as described above. Following which RNA was extracted using TRI reagent (Sigma-Aldrich) using protocol provided with the reagent. The RNA was converted into cDNA using Verso cDNA synthesis kit (Thermo Fisher Scientific). Two μL of this cDNA was mixed with 10 μL of 2x qPCR mix Radiant™ SYBR Green Lo-ROX (Alkali Scientific), 1 μL of 10 μM forward (Fwd) primer, 1 μL of 10 μM reverse (Rev) primer and 6 μL of nuclease free water. PCR settings were 50°C for 2 min, 95°C for 10 min, then 40 cycles at 95°C for 15 sec and 60°C for 1 min. Following PCR completion, melting curve was recorded using these settings: 95°C for 15 sec, 60°C for 1 min and 95°C for 15 sec.

Sequences of the primers used for RT-PCR:

SMOX Fwd 5’ TCAAAGACAGCGCCCAT 3’; SMOX Rev 5’ CCGTGGGTGGTGGAATAGTA 3’

PAOX Fwd 5’ACTAGGGGGTCCTACAGCTA 3’; PAOX Rev 5 ‘CGTGGAGTAAAACGTGCGAT 3’

### Metabolite extraction

Retinal endothelial cells were plated in 100 mm dishes coated with attachment factor (Cell Systems) at density of 0.9 or 0.4 x 10^6^ cells per plate and maintained in endothelial cell media (CellBiologics) in a 5% CO_2_ incubator set at 37°C for 3 days. After 3 days of incubation, media was changed to high glucose DMEM media (Cleveland Clinic Media lab) without FBS and cells were again incubated in normoxic incubator, to synchronize the cells. After 6h, media was changed back to endothelial cell media (Cell Biologics) and plates were either incubated in normoxic or hyperoxic (75% oxygen), 5% C0_2_ incubator set at 37°C for 24 h. Following 24 h of incubation, metabolites were extracted. To extract metabolites, media was aspirated, plates were washed with 10 ml of room temperature normal saline. To the washed cells 300 μL of 0.1% formic acid (prepared in water) containing 1 μg of ^13^C_5_ ribitol per ml of solution. Next, 600 μL of −20°C cold methanol was added to each plate. Cells were scraped with a cells scraper while keeping plates on ice and cell lysates were transferred to tubes containing 450 μL of −20°C cold chloroform. Tubes were agitated on a thermomixer at 4°C, 1400 rpm for 30 min. Tubes were then centrifuged for 5 min at 15000 x g at 4°C. Six-hundred microliters of supernatant was transferred to fresh tubes and dried under vacuum in a −4°C cold vacuum evaporator (Labconco). Samples were derivatized with the two step protocol as described earlier (Singh et. al. 2020) and measured using GCMS method described earlier (Singh et. al. 2020).

## Acknowledgements

Grant Support: National Eye Institute (R01 EY024972 to JES; P30 EY025585 to Ophthalmic Research); Research to Prevent Blindness Physician Scientist (RPB1801 to JES).

