## Supplemental Data for "Hyperoxia inhibits proliferation of retinal endothelial cells in Myc dependent manner"

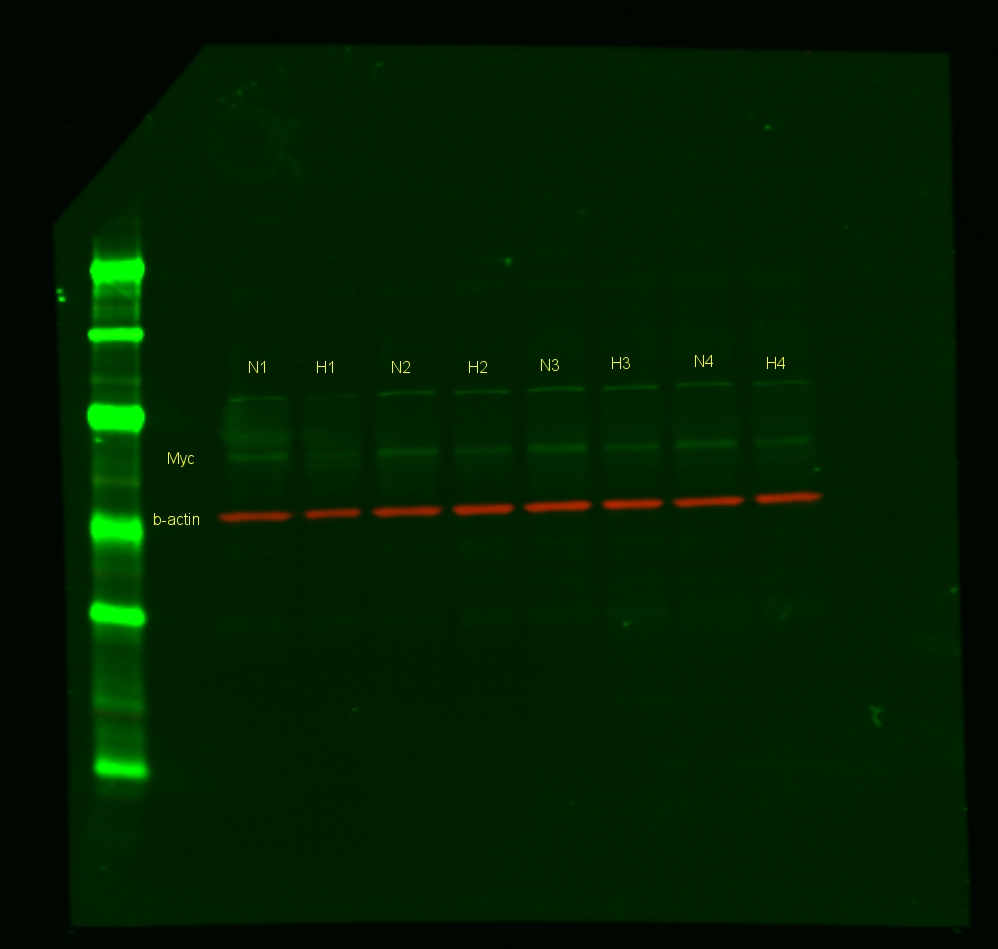

**Figure S1 Confirmatory Western blot analysis of Myc** **protein.** N1-4, normoxic samples; H1-4, hyperoxic samples (n=4 biological replicates per condition)

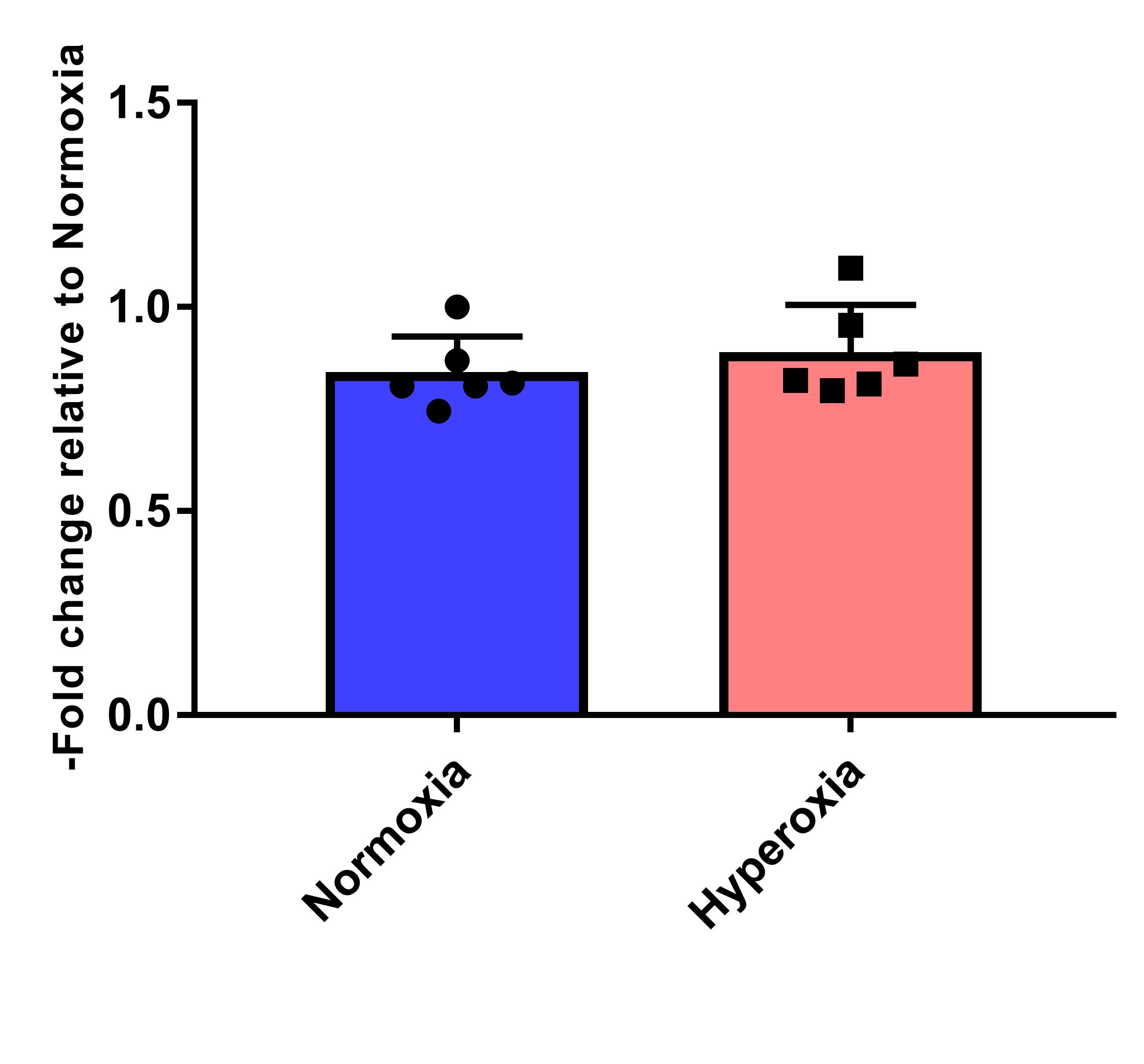

**Figure S2 Expression levels of cMyc in the retinal endothelial incubated in normoxic or hyperoxic conditions** (n=6 biological replicates per condition).

**a)**

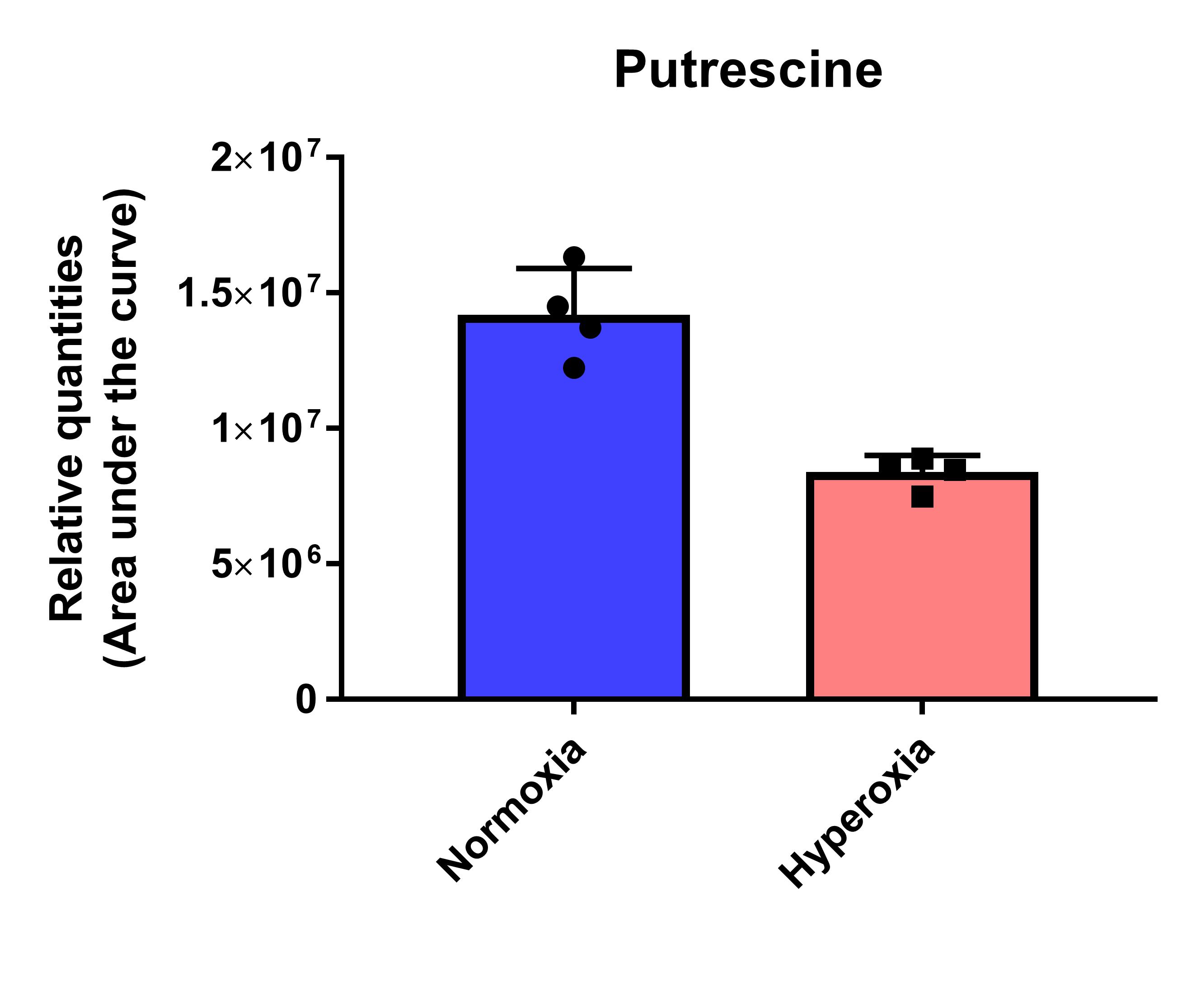

**b)**

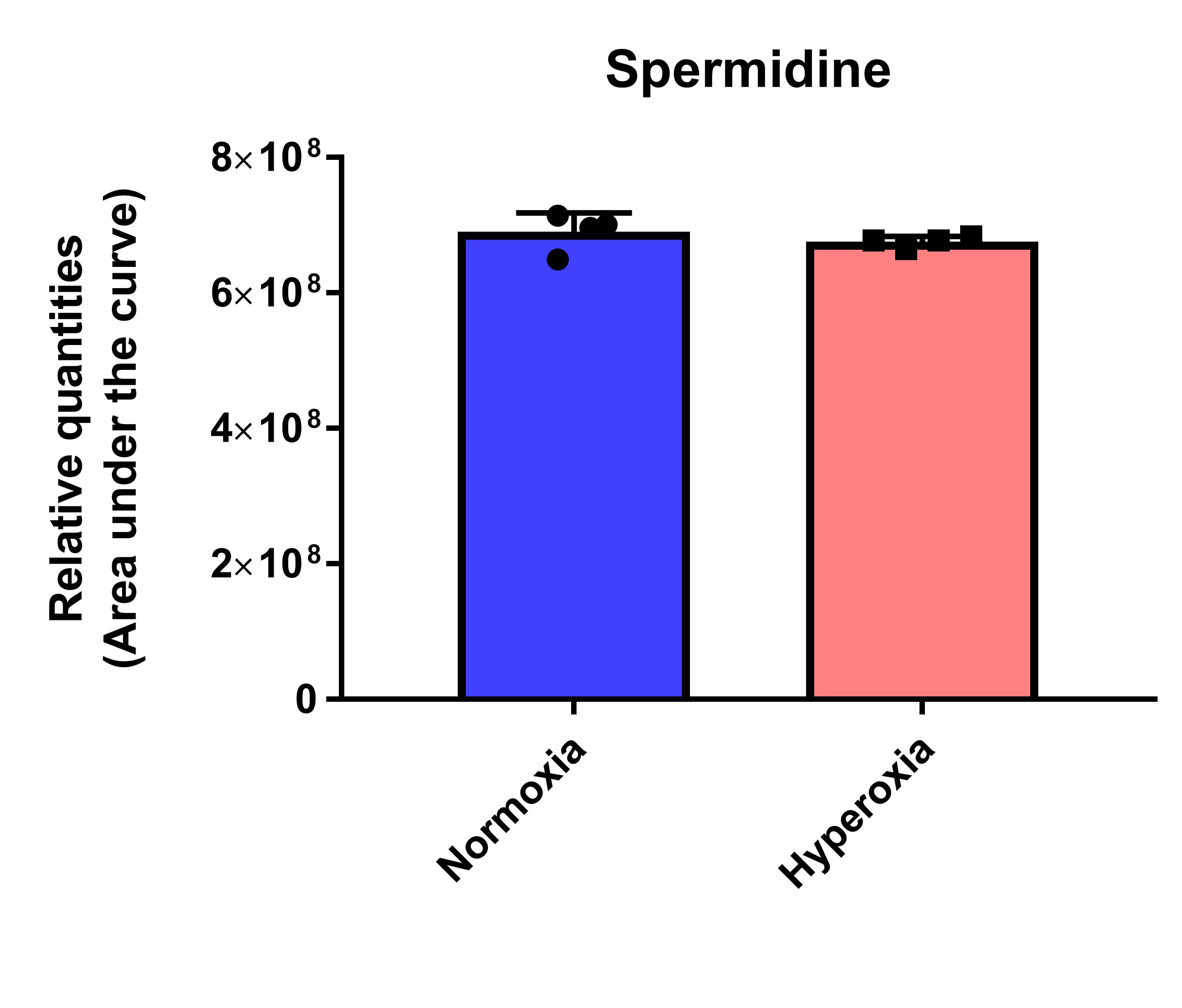

**Figure S3 Hyperoxia leads to decreased levels of polyamine pathway precursor putrescine.** a) Relative quantities of putrescine quantified as area under the curve for XIC m/z 89.10732 [M+H]^+^, and b) spermidine XIC m/z 146.16517 [M+H]^+^. (n=4 biological replicates per condition). t-test p-values, <0.05 for putrescine and non-significant for spermidine.

**a)**

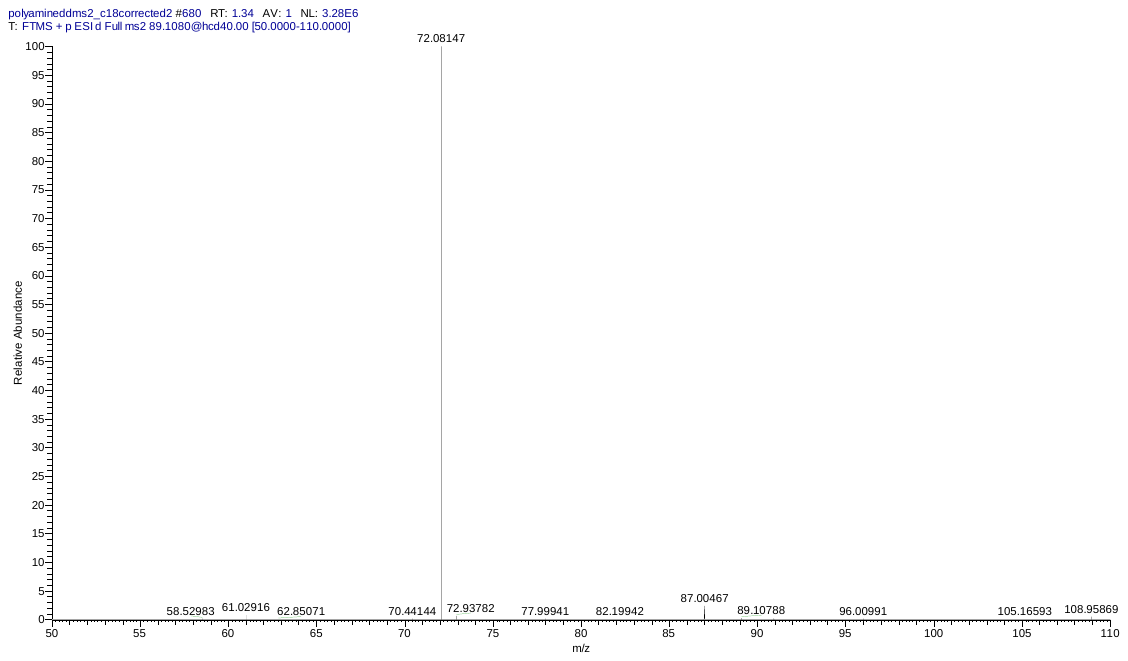

**b)**

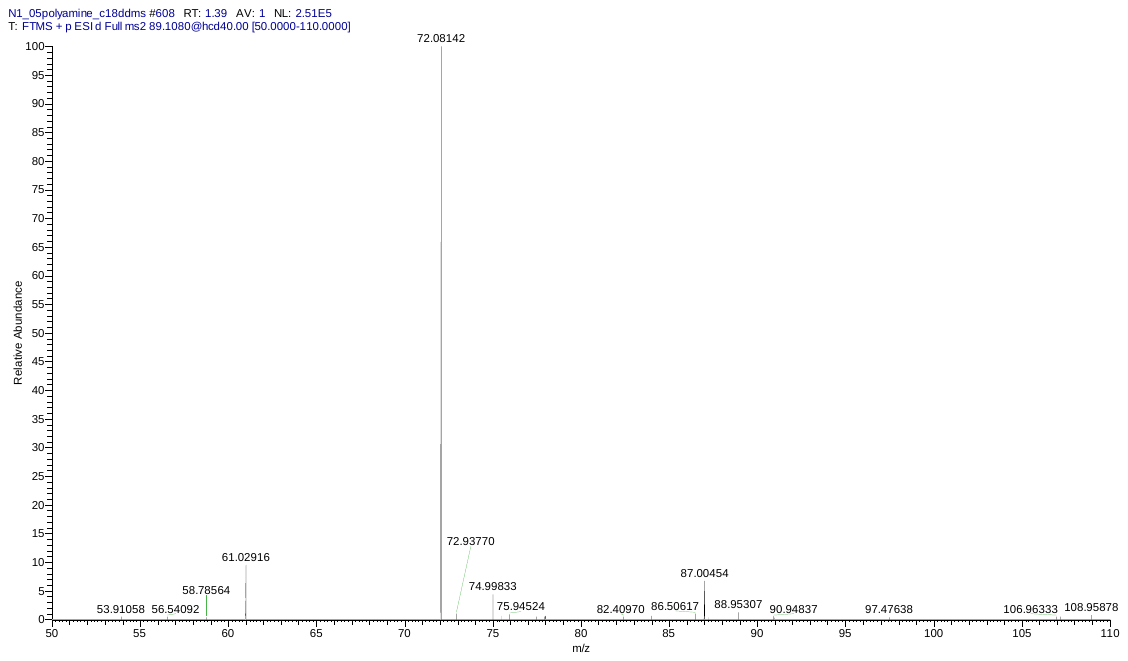

**Figure S4 MS2 fragmentation of putrescine (m/z 89.1073, [M+H]^+^)** in a) authentic standard and b) biological sample

**SUPPLEMENTAL METHODS**

**SDS-PAGE and Western blotting**

SDS-PAGE and western blotting was performed as described in the main manuscript with exceptions described here. Sample 25 μL of supernatant per sample was loaded into each well of 4-20% Tris-glycine Novex^TM^ WedgeWell^TM^ precast gel (Invitrogen). Samples were separated at a constant voltage of 190 V. Steps following this step were the same as described in the main manuscript. The antibodies used in this Myc confirmatory blot were: c-Myc (E5Q6W) Rabbit mAb #18583 and β-actin (8H10D10) Mouse mAb catalog # 3700 (Cell signaling). With this cMyc antibody, the blot was not stripped/re-probed.

**LC-MS/MS measurements of polyamines**

Metabolites were separated on Pursuit XRs C18 (150 × 2.0 mm, 5 μm, 100 Å; Agilent). The LC method used was similar to that published earlier (Singh et al., 2017). Briefly, a gradient of solvent A (0.1% formic acid in water) and solvent B (0.08% formic acid in acetonitrile) was ramped from 10% B to 80% B in 4 minutes, with a wash step with 80% B from 7 to 8 minutes and re-equilibration with 10% B from 8 to 10 minutes. Solvent flow rate was set to 200 μL/min and the column oven was maintained at 30°C. The autosampler was maintained at 4°C and 5 μL of sample was injected. Heated electrospray ionization (ESI) source settings used for the mass spectrometer were spray voltage 3500 in positive mode, capillary temperature 250°C, sweep gas flow rate of 2, sheath gas 45, S-lense RF 50, and auxiliary gas 10.

The mass spectrometer method used was data-dependent MS2, where a full scan (survey) was recorded followed by MS2 scans for the top 3 peaks seen in the survey full scan. The inclusion list contained m/z values 89.10732, 146.16517, 203.22302 and with 10 ppm mass error. MS1 parameters were MS1 m/z window set from 50 to 750 mass cutoff, resolution of 120,000, automatic gain control of 3 × 10^6^ and maximum injection time of 100 ms. MS2 values were recorded for the top 3 peaks after every MS1 scan, with 30,000 mass resolution, an automated gain control of 10^5^, maximum injection time of 50 ms, and precursor ion isolation width of 1 m/z. A normalized HCD collision energy of 40% was applied. To exclude the redundant MS2 scans a dynamic exclusion of 10 seconds was applied. Isotope exclusion was applied to reduce dwell time for unnecessary MS2 scans.
